# Multiscale Co-Expression in the Brain

**DOI:** 10.1101/2020.03.31.018630

**Authors:** Benjamin D. Harris, Megan Crow, Stephan Fischer, Jesse Gillis

**Affiliations:** Cold Spring Harbor Laboratory, Stanley Institute for Cognitive Genomics, Cold Spring Harbor, NY, USA; Cold Spring Harbor Laboratory, Watson School of Biological Sciences, Cold Spring Harbor, NY, USA

## Abstract

Single-cell RNA-sequencing (scRNAseq) data can reveal co-regulatory relationships between genes that may be hidden in bulk RNAseq due to cell type confounding. Using the primary motor cortex data from the Brain Initiative Cell Census Network (BICCN), we study cell type specific co-expression across 500,000 cells. Surprisingly, we find that the same gene-gene relationships that differentiate cell types are evident at finer and broader scales, suggesting a consistent multiscale regulatory landscape.

## Main

The seven mouse primary motor cortex scRNAseq datasets, totaling over 500,000 cells/nuclei, from the Brain Initiative Cell Census Network provide the first opportunity to comprehensively study cell type specific co-expression networks with scRNAseq data (Figure 1A)^1^. This is important since previous work has shown that compositional differences confound co-regulatory signals in co-expression networks generated from bulk expression data^2,3^. Though our previous research suggested that single-cell data showed similar co-expression to bulk to a surprising degree^4^, further analysis in more specific and powered data will advance our understanding of both regulatory and cell type specific co-expression signals. In the BICCN datasets, replicable annotations of cell types at multiple levels of classification enable meta-analysis of cell type specific co-regulatory modules (Supplementary Figure 1).

**Figure 1.**
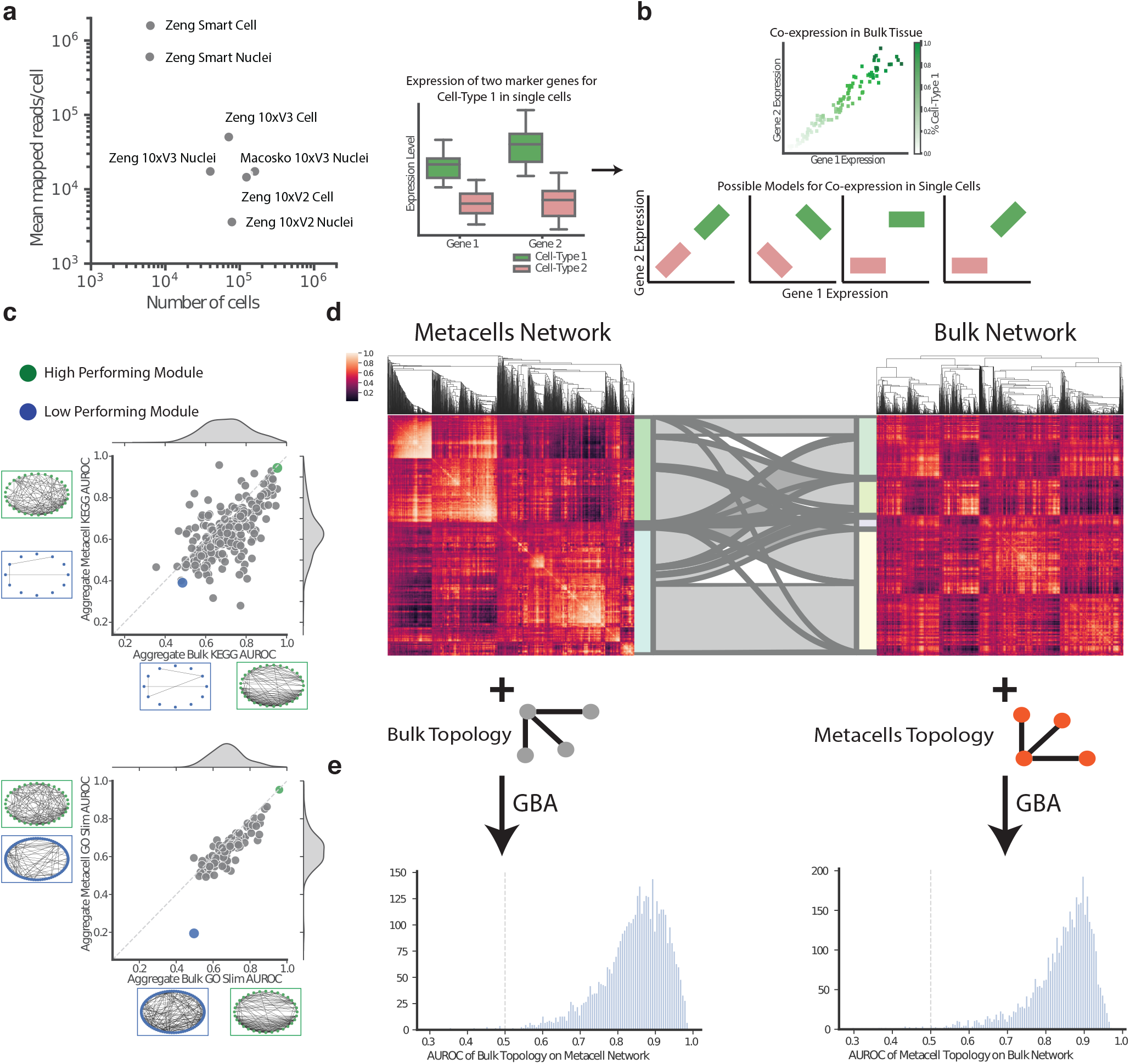
Co-regulatory and compositional co-expression networks share topology and functional connectivity. **a.** The single cell datasets were generated with multiple technologies, resulting in varying numbers of cells and varying read depth across datasets. **b.** In bulk RNAseq data, marker genes must be co-expressed because of compositional differences in samples. Co-expression within individual cell types can take multiple different models. **c.** Performance of KEGG and GO on both metacells and bulk RNAseq co-expression networks is well correlated. Network diagrams show the best performing (Green) and worst performing (Blue) terms in each dataset. **d.** Clustered heatmaps for bulk metacells and bulk networks. The riverplot joining them identifies shared genes across hierarchical clusters. **e.** Prediction of small neighborhoods in one network using the other network’s topology shows shared local topology.

Our basic strategy is to build networks based on the known hierarchy of cell types within the BICCN data, and to evaluate the co-expression of cell type markers in networks that control for this source of variation. Thus, for example, two genes which are highly expressed in cell type A relative to cell type B will be co-expressed in a network containing both cell types since the genes are co-variable with respect to cell type. The fundamental question of single-cell co-expression is the degree to which novel covariation is present in cell type A (or B) individually, reflecting regulatory interactions rather than compositional effects (Figure 1B)^5^. We use meta-analysis to identify robust edges (Supplementary Figure 2), and define “co-regulatory” vs. “compositional” gene interactions by building networks which control for differing levels of cell type variation. At the finest level are metacell networks which measure gene-gene co-variation over statistically similar sets of cells^6^. By comparing gene-gene relationships that sample from more and more diverse cells, we begin to incorporate compositional effects across the types of cells sampled (e.g., subtypes of inhibitory cells). At the broadest level of analysis are bulk co-expression networks, using samples made up of large numbers of cells^7^.

For our first experiment, we compared metacell networks to bulk in order to capture similarities and differences at the farthest range of the spectrum (see Methods for details on network construction). We first establish that both networks reflect known biology using a guilt-by-association formalism, in which each network is measured for its ability to reconstruct a partially hidden gene list from preferential connectivity within it, outputting an Area under the ROC curve (AUROC) (Figure 1B, Supplementary Figure 3)^8^. In the aggregate metacells network, the average AUROC for GO slim and KEGG are 0.64 and 0.63 respectively, and similarly the average AUROC of the aggregate bulk RNAseq network is 0.67 for GO and 0.70 for KEGG (Figure 1C). We also find that these networks have highly similar topologies. A comparison of coarse hierarchical clustering of both co-expression networks shows large shared modules between the two networks, visualized as a riverplot in Figure 1D. Moreover, the average AUROC of modules drawn from the metacell network in the bulk network is 0.84, and the same is true of the reverse analysis (Figure 1E). This indicates that modules present in one are present in the other to a very specific degree, which is surprising since these two networks were constructed using data that capture vastly different signals, compositional versus co-regulatory.

To investigate the overlap between compositional and co-regulatory variation more directly, we next evaluated the modularity of neuronal subclass markers in each of these two networks. As expected, the markers are well connected in meta-analytic networks built from bulk RNA-seq (average AUROC=0.84, Figure 2A), consistent with the notion that these networks contain cell type signals. Surprisingly, markers are also well connected in the networks where cell type variation has been controlled (average AUROC=0.84, Figure 2B). The performance of the subclass markers in both networks is well correlated (r=0.73, p=0.004, Figure 2C), in agreement with the consistent topology we find in both networks. This suggests that whatever regulatory factors drive differences between cell types remain important as a source of differences within cell types.

**Figure 2.**
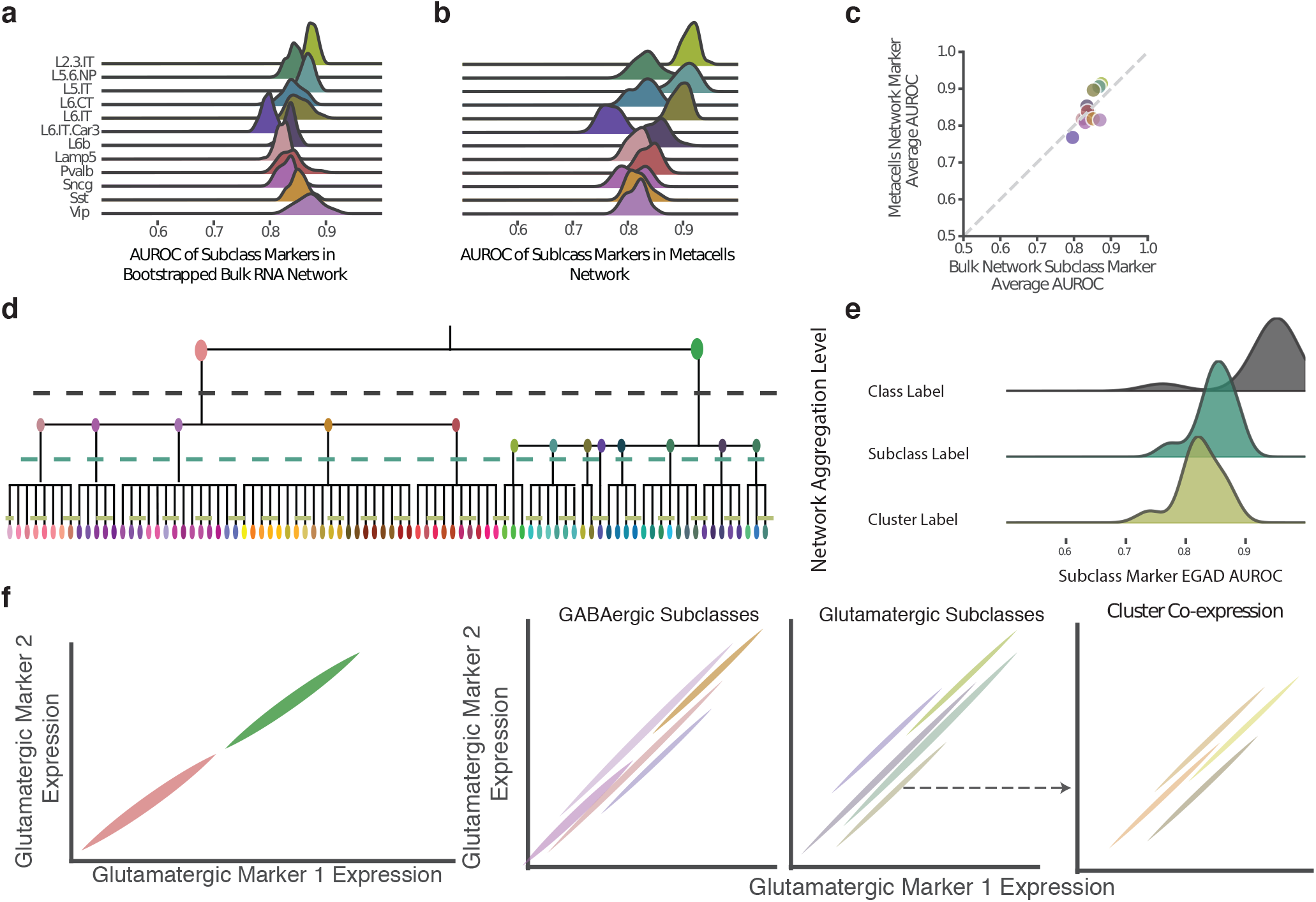
Cell type marker sets show multiscale co-expression. **a.** Markers show high performance in bulk RNAseq networks. **b.** Subclass markers show consistent and high co-expression in metacell networks. **c.** Average performance of bulk and metacell networks is correlated. **d.** Dendrogram of cell type hierarchy showing class, subclass, and clusters used to construct co-expression networks. **e.** Consistent and strong co-expression of markers in networks at each level of the cell type hierarchy. **f.** A multiscale model for co-expression of a pair of glutamatergic markers shows co-expression of the markers through all levels of the hierarchy of cell types.

The BICCN data affords the unique opportunity to use consistent cell type labels across independently sampled datasets, so that robust analyses can be constructed at varying levels of specificity in the cell type hierarchy across independent data. We took advantage of the known hierarchy for our next series of experiments. For each of three levels of classification, class, subclass and cluster, we build aggregate networks to capture replicable gene-gene relationships (Figure 2D). When we evaluate subclass level markers, the class network will be compositional, but the subclass and cluster networks will be non-compositional. While the class network has the highest performance (average AUROC=0.94), we find the subclass and cluster networks still perform exceptionally well (subclass: average AUROC=0.85, cluster: average AUROC=0.83, Figure 2E). This is also true of subsampled networks that reduce within-cluster heterogeneity, further strengthening this observation (Supplementary Figure 4). These results indicate that similar gene-gene interactions occur both within and across cell types (i.e., co-expression is multiscale).

To visualize this surprising result, we focus on changes in networks for two of the GABAergic subclasses: Vip and Sst. In the class network, the subclass markers are extremely modular, with dense connectivity within each gene list and sparse connections between them. However, for the subclass and cluster networks, the connections between the modules increase significantly. Despite this increase, the Vip and Sst modules can still be clearly discerned from each other. We quantify the change in connectivity between modules by measuring how one gene list, the training list, predicts another gene list, the testing list. As expected from the network diagrams, the AUROCs indicate higher connectivity between the Vip and Sst modules in the subclass and cluster level networks (Supplementary Figure 5). Moving past modules of genes to individual pairs, we continue to see an important role for multiscale co-expression (Supplementary Figure 6).

The shared co-expression signal of the marker genes and regulatory modules throughout the cell type hierarchy makes it clear that the co-expression is, in part, multiscale. We illustrate the persistent co-expression relationship of marker genes at all levels of cell type classification using a pair of hypothetical glutamatergic markers (Figure 2F). The multiscale co-expression shows that, while gene expression values are significantly different at high levels of classification, the core co-regulatory network remains constant throughout the cell type hierarchy.

## Methods

### Single Cell Datasets and Preprocessing

We acquired the datasets and associated metadata directly from the BICCN. To work with the expression data, anndata objects were created by filtering the droplets to the whitelist defined by the consortium and merging with all associated metadata. All analyses were done using CPM normalized expression values. To select a shared list of genes we ranked each gene by its average expression and selected the top 7,500 genes in each dataset. Then genes that were in the top 7,500 for at least 6 of the 7 datasets were used in all analysis, leaving us with 4,201 genes. All analyses were done with this list of genes.

### Bulk RNA Sequencing Data from GEMMA

Metadata from the Gemma database^7^ was acquired on 11-29-19. The metadata was filtered to include only mouse bulk RNAseq datasets with at least 20 samples. Then metadata terms were filtered for relevance for the brain, leaving 29 terms (see github repository). The expression data was then downloaded using the GEMMA R API and filtered to the same genes as the scRNAseq data. Networks were built as detailed below.

### Network Construction and Aggregation

Networks were built by rank standardizing the Pearson correlation matrix of the genes. After ranking, we replace the undefined values with the average of the network. For the bulk data networks are built using an entire dataset. For single cell datasets, a full compositional network is computed using only the labeled neurons in each dataset. When computing class, subclass, cluster, and metacell networks we partition each dataset by the metadata label and build a network for each value. After aggregating networks within each dataset, we aggregate the ranked dataset networks. Aggregating datasets occurs by summing the networks from each dataset and then ranking the sum.

When computing network performance of markers in the bulk network we bootstrap the bulk datasets 100 times to create 100 networks. In the down-sampling experiment we compute centroids for each dataset partition and select the 50 closest cells to the centroid within that partition. We exclude any partition with fewer than 100 cells to make sure it is at most 50% of the original data in the partition.

### Computing Marker Genes

Marker gene lists are computed using the Mann-Whitney test in each dataset using a 1vsAll design. Significance is computed with a threshold of log2FC >2 and FDR <0.05. To compute markers across datasets we compute recurrence of each gene by totaling the number of datasets the gene is significantly different in. After sorting genes by recurrence, we sort by average AUROC. We used gene sets of size 100. Subclass-specific markers are computed within classes, e.g. Vip markers are extracted by finding genes that are differentially expressed with respect to all other GABAergic subclasses. For example when computing the markers for the Vip subclass, a GABAergic subclass, we only compare the expression of the Vip cells to the other GABAergic subclasses.

### Measuring Network Performance with EGAD

A python version of the R package EGAD was created by translating the runGBA() function from the R package. It was modified to do cross validation in known splits, instead of randomly partitioning the data. We run it with 3 fold cross-validation. The algorithm uses neighbor voting to compute the sum of ranks of predictions for a given gene set within a network. Using the sum of predicted ranks we calculate an AUROC and/or a p-value as an output.

### Computing Metacells

Metacells are computed using the R metacells package. We set the parameters to encourage extremely small clusters (K=20, m=5, b=1000). Additionally, we used the 4,201 recurrently highly expressed genes as the gene list for the method. While the metacells method, like the original clustering method, is graph based, minor differences in the methods allow for metacell clusters that contain multiple subclasses. To avoid any compositional effects, we filter out all metacell clusters containing cells from multiple subclasses.

### Code Availability

We deposit the code used in www.github.com/bharris12/multiscale_brain. The code includes core functions for the different algorithms used as well as the code used to create the individual figures.

## Acknowledgements

BH was supported by the CSHL Crick Cray Fellowship. MC was supported by NIH K99MH120050. JG and SF were supported by NIH R01MH113005, R01LM012736 and U19MH114821.

## Author Contributions statement

JG conceived the study. JG, MC and BH conceived the experiments. BH and SF performed the computational analysis. All authors read and approved the final manuscript.

## Additional Information

See Supplementary Information for Supplementary Note 1 and Supplementary Figures

## Notes

https://github.com/bharris12/multiscale_brain

## References

1. Yao, Z. et al. Biorxiv 2020.02.29.970558 (2020).doi:10.1101/2020.02.29.970558

2. Farahbod, M. Pavlidis, P. Biorxiv 735951 (2019).doi:10.1101/735951

3. Zhang, Y., Cuerdo, J., Halushka, M.K. McCall, M.N. Brief Bioinform (2019).doi:10.1093/bib/bbz135

4. Crow, M., Paul, A., Ballouz, S., Huang, Z.J. Gillis, J. Genome Biol 17, 101 (2016). doi: 10.1186/s13059-016-0964-6

5. Trapnell, C. Genome Res 25, 1491–1498 (2015). doi: 10.1101/gr.190595.115

6. Baran, Y. et al. Genome Biol 20, 206 (2019). doi: 10.1186/s13059-019-1812-2

7. Zoubarev, A. et al. Bioinform 28, 2272–3 (2012). doi: 10.1093/bioinformatics/bts430

8. Ballouz, S., Weber, M., Pavlidis, P. Gillis, J. Bioinformatics btw695 (2016).doi:10.1093/bioinformatics/btw695

