## Supplementary Information for "Multiscale Co-Expression in the Brain"

March 31, 2020

1. Cold Spring Harbor Laboratory, Stanley Institute for Cognitive Genomics, Cold Spring Harbor, NY, USA 2.Cold Spring Harbor Laboratory, Watson School of Biological Sciences, Cold Spring Harbor, NY, USA

#### Contents

1 Supplementary Note 1: Connectivity Between Subclass Modules in Non-Compositional Co-expression Networks 1

2 Software Used 2

3 Supplementary Figures 2

#### 1 Supplementary Note 1: Connectivity Between Subclass Modules in Non-Compositional Co-expression Networks

We examine the change in module connectivity for all subclasses by comparing cross module performance for random modules predicting random modules, markers predicting markers and random modules predicting markers using all subclasses and 200 random gene sets (Supplementary Figure 5). Pairs of random modules have random connectivity between each other in networks at all levels of the hierarchy: class, subclass and cluster level (Avg AUROC = .49). In the class level network, subclass modules also have random connectivity with other subclass modules (Avg AUROC .50). However, the Avg AUROC for subclass and cluster level networks is .65 and .66 respectively. The significant increase in average performance of marker sets predicting other marker sets matches the trend we find with the Vip and Sst modules. Finally, random modules all predict subclass markers with an Avg AUROC of .49. However, the distributions have much more variance than the null model. We find that performance is correlated with the percent of random modules that are genes within a subclass marker list (Class: r=.32, p=3.9e-6; Subclass r=.42, p=6.06e-10, Cluster=.4, p=2.9e-9; Supplementary Figure 5). Through this analysis, we show consistent modules of co-expression for subclass level markers, with increasing inter-module connectivity in non-compositional networks. These results indicate that core co-regulatory relationships span cell-type marker gene lists, while marker gene lists even though marker gene lists show strong co-regulation on their own.

### 2 Software Used

The main software packages used in the analysis: Scanpy1, Scipy2, Matplotlib3, Seaborn, Bottleneck, GNU Parallel4, Metacells5, networkx6, scrattch7, Pandas, Numpy8 Joypy9. We have a detailed list of versions and software in the github repository.


1. Wolf, F. A., Angerer, P. Theis, F. J. SCANPY: large-scale single-cell gene expression data analysis. Genome Biol 19, 15 (2018).

2. Virtanen, P. et al. SciPy 1.0: fundamental algorithms for scientific computing in Python. Nat Methods 17, 261–272 (2020).

3. Hunter, J. D. Matplotlib: A 2D Graphics Environment. Comput Sci Eng 9, 90–95 (2007).

4. Tange, O. GNU Parallel 20150322 ('Hellwig'). (2015) doi:10.5281/zenodo.16303.

5. Baran, Y. et al. MetaCell: analysis of single-cell RNA-seq data using K-nn graph partitions. Genome Biol 20, 206 (2019).

6. Hagberg, A. A., Schult, D. A. Swart, P. J. Exploring Network Structure, Dynamics, and Function using NetworkX. in Proceedings of hte 7th Python in Science Conference (2008).

7. Tasic, B. et al. Adult mouse cortical cell taxonomy revealed by single cell transcriptomics. Nat Neurosci 19, 335–346 (2016).

8. Walt, S. van der, Colbert, S. C. Varoquaux, G. The NumPy Array: A Structure for Efficient Numerical Computation. Comput Sci Eng 13, 22–30 (2011).

9. Taccari L. Joypy https://github.com/sbebo/joypy


### 3 Supplementary Figures

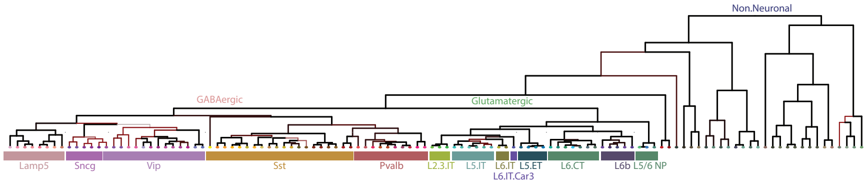


Supplementary Figure 1: **Dendrogram of cell-type hierarchy in scRNAseq datasets**

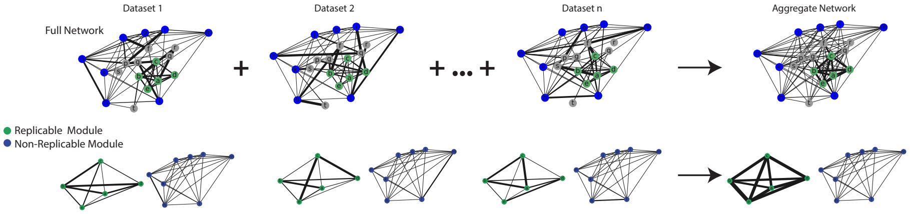


Supplementary Figure 2: **Meta-analysis across dataset networks identifies robust co-expression relationships**

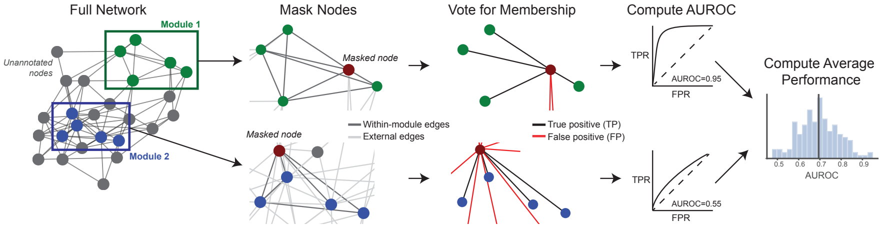


Supplementary Figure 3: **Guilt-By-Association algorithm assess a network's ability to reconstruct modules**

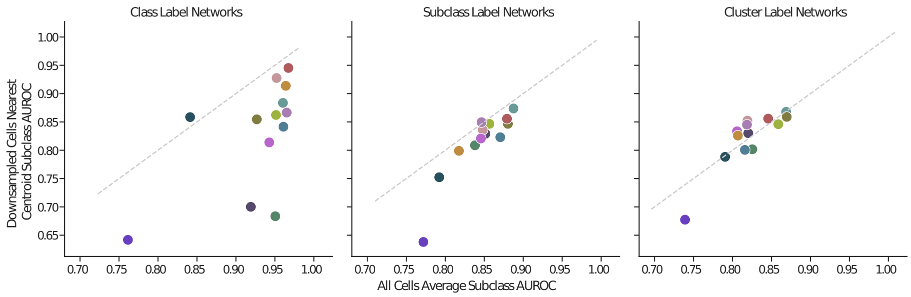


Supplementary Figure 4: **Subsampled datasets reconstruct subclass modules as well as full data** Comparison of performance of subclass level markers using full datasets and downsampling of each cluster to the 50 cells nearest the centroid of the cluster. Coloring matches subclass coloring in Supplementary Figure 1 dendrogram.

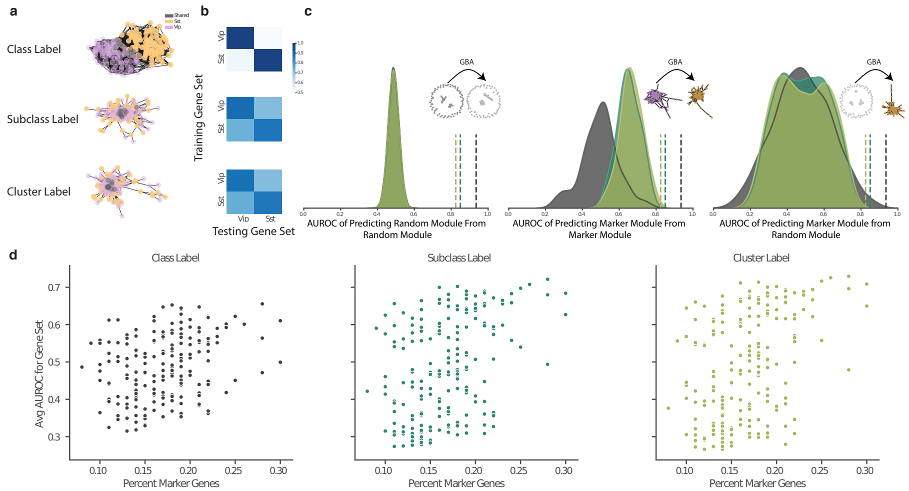


Supplementary Figure 5: **Stronger connectivity between subclass modules in non-compositional networks. a.** Network diagrams show strong connectivity within both Vip and Sst modules at all levels and increased connectivity between the modules at the subclass and cluster level networks. **b.** Predicting connectivity between modules show random connections at the class level and strong connections at the subclass and cluster level for Vip and Sst modules. textbfc. Random gene sets have random connectivity to other random gene sets in all networks, while subclasses have non-random connectivity between each other in only subclass and cluster level networks. Random gene sets have variable levels of connectivity to subclass modules **d.** Random gene set connectivity to subclass modules is correlated to the percent of random genes that are a marker in any subclass.

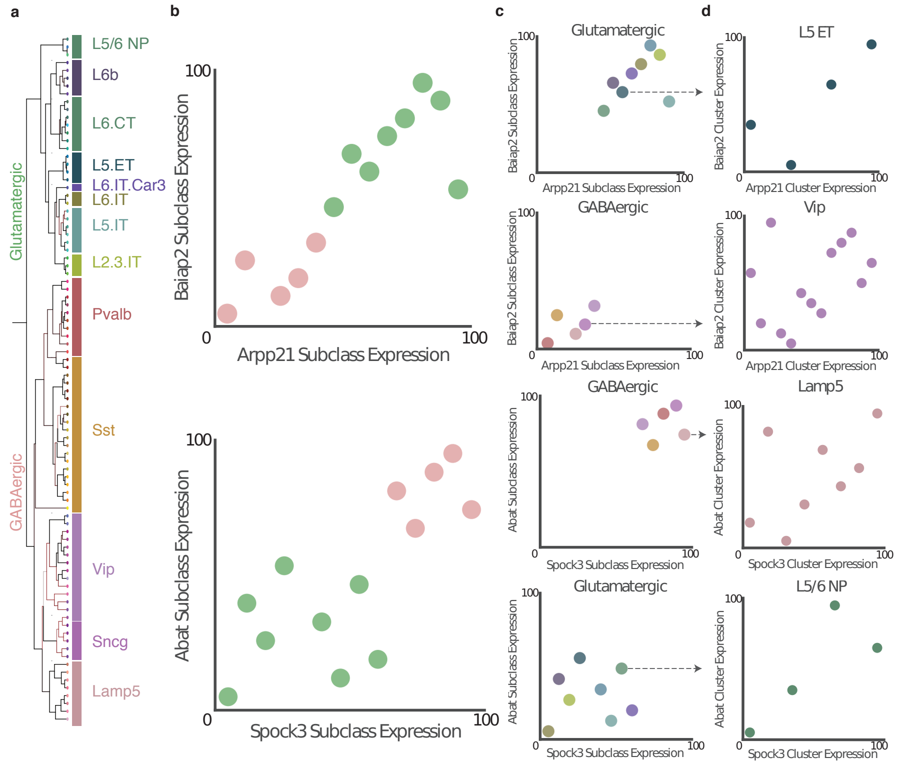


Supplementary Figure 6: **Multi-scale co-expression in pairs of marker genes. a.** A dendrogram of cell-type hierarchy and colors. **b.** Expression percentiles aggregated across datasets for a pair of Glutamatergic markers, Arpp21 and Baiap2, and GABAErgic marker, Spock3 and Abat. **c.** The GABAergic and Glutamatergic markers remain co-expressed when split into GABAergic and Glutamatergic subclasses. **d.** Within subclasses clusters are co-expressed in both GABAergic and Glutatmatergic subclasses.

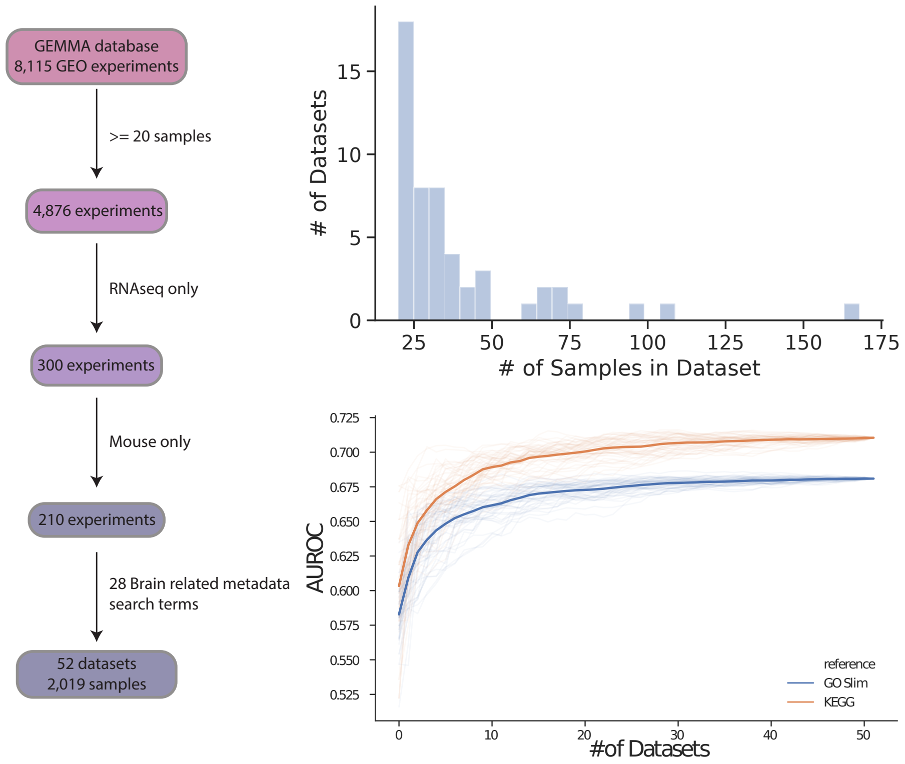


Supplementary Figure 7: **Co-expression of bulk RNAseq data from the brain**

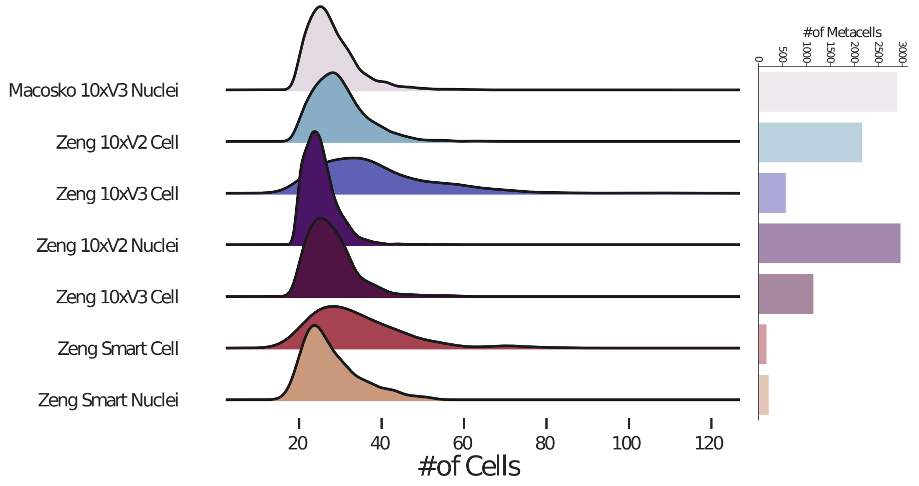


Supplementary Figure 8: **Number and size of metacells computed from scRNAseq datasets**
